# The ACE-2 receptor accelerates but is not biochemically required for SARS-CoV-2 membrane fusion

**DOI:** 10.1101/2022.10.22.513347

**Authors:** Marcos Cervantes, Tobin Hess, Giorgio G. Morbioli, Anjali Sengar, Peter M. Kasson

## Abstract

The SARS-CoV-2 coronavirus infects human cells via the ACE-2 receptor. Circumstantial evidence suggests that ACE-2 may not just serve as an attachment factor but also help activate the SARS-CoV-2 spike protein for membrane fusion. Here, we test that hypothesis directly, using DNA-lipid tethering as a synthetic attachment factor in the place of ACE-2. We find that SARS-CoV-2 pseudovirus and viruslike particles are both capable of membrane fusion if attached in the absence of ACE-2 and activated with an appropriate protease. However, addition of soluble ACE-2 speeds the fusion reaction. This is observed for both the Wuhan strain and the B.1.1.529 Omicron variant. Kinetic analysis suggests that there are at least two rate-limiting steps for SARS-CoV-2 membrane fusion, one of which is ACE-2 dependent and one of which is not. These data establish that, in the presence of an alternative attachment factor, ACE-2 is not biochemically required for SARS-CoV-2 membrane fusion. Since ACE-2 serves as the high-affinity attachment factor on human cells, the possibility to replace it with other factors has implications for the evolvability of SARS-CoV-2 and the fitness landscape for future related coronaviruses.

## INTRODUCTION

SARS-CoV-2 has caused a global pandemic since its emergence in humans, with over 600 million confirmed infections and 6.5 million deaths as of October 2022^1^. Viral entry and infection are mediated by the SARS-CoV-2 spike protein, which binds to ACE-2 receptors on the cell surface^2,3^, is activated via proteolytic cleavage^4^, and then drives membrane fusion between the viral envelope and a cellular membrane. Since the spike protein performs multiple roles, dissecting the functional requirements of each can be challenging. Separating out these requirements, however, will yield an understanding not only of how SARS-CoV-2 currently infects cells but a better ability to predict future evolution of this and similar viruses.

The SARS-CoV-2 spike protein binds to ACE-2 via its receptor-binding domain (RBD). Structural studies have identified “down” and “up” conformations of the RBD in the spike trimer, with “up” conformations capable of binding ACE-2. Single-molecule FRET experiments have also shown that the RBD occupies a conformational equilibrium that is modulated by ACE-2. Further computational studies have postulated a broader range of “down” and “up” conformations^5,6^, and additional structures of SARS-CoV-2 spike in complex with ACE-2 have suggested that ACE-2 binding can destabilize the trimer^7^. This destabilization is believed important to activation for membrane fusion, which requires proteolysis at the S2’ site on the spike protein^7–9^. Such proteolysis can be performed by a number of enzymes, most notably TMPRSS2 on the cell surface, but also endosomal cathepsins and extracellular proteases^10–13^.

These observations naturally yield a model where ACE-2 binding primes SARS-CoV-2 spike for proteolysis, release of the fusion peptides, and ultimately membrane fusion and entry. Such receptor-activated fusion has been observed in many strains of HIV, where receptor/co-receptor ligation is critical for conformational activation of the envelope protein^14–17^. In contrast, receptor binding by influenza hemagglutinin appears primarily to localize the virus near the plasma membrane, since replacement of physiological receptors with synthetic tethers anchored in the viral membrane can functionally reconstitute fusion with identical kinetics^18^. Here, we ask where SARS-CoV-2 falls upon this continuum in the requirement of receptor binding for conformational activation of the spike protein.

To accomplish this, we use DNA-lipid tethers that can irreversibly insert into membranes and permit programmable self-assembly.^19^ Complementary strands of DNA conjugated to lipids inserted in viral particles and synthetic liposomes can thus target the particle to the liposome in the absence of physiological receptors. The triggers for fusion can then be chemically reconstituted and tested precisely. Fusion can be detected using single-virus optical microscopy, where fluorescent dyes in the virus or the target membrane report on viral state changes occurring through the fusion process^20,21^. This approach of DNA-lipid tethering and single-virus fusion has previously been used to characterize entry by multiple different viral families, including orthomyxoviruses^18^ and flaviviruses ^22^.

Separating receptor binding from membrane fusion permits examination of the biochemical requirements for viral entry, which may be distinct from the most common pathways a given virus takes for infection. There has been extensive work on the cellular pathways for SARS-CoV-2 entry^12,23–27^. As the evolution of SARS-CoV-2 has shown, however, the most common entry pathways of a virus can change even over the space of a few years--the Wuhan strain primarily entered via TMPRSS2-mediated fusion, whereas the BA.1 strain primarily entered via cathepsin-mediated fusion^25,26^. Understanding the biochemical requirements for entry thus provides critical knowledge to help predict future evolutionary pathways of SARS-CoV-2 and their respective vulnerability to therapeutic countermeasures.

## MATERIALS AND METHODS

Full experimental details are given in the Supporting Information. Briefly, HIV pseudoviruses were produced using previously published protocols ^28^ and plasmids that were gifts of Jesse Bloom. Omicron spike pseudotyping was performed using the plasmid pTwist-SARS-CoV-2 Δ18 B.1.1.529, a gift from Alejandro Balazs ^29^. Virus-like particles were produced using plasmids encoding the M, N, E, and S proteins from SARS-CoV-2 in addition to a luciferase RNA carrying the SARS-CoV-2 PS9 sequence using previously published protocols ^30^. Plasmids were a gift from Jennifer Doudna. Viral particles were labeled with Texas Red-DHPE at a quenching concentration. All handling of pseudoviral and virus-like particles was performed under BSL-2 conditions using institutionally approved protocols.

Target liposomes were composed of 68.75 mol% POPC, 20% DOPE, 10 % cholesterol, 1% biotin-DPPE, 0.25 % Oregon Green-DHPE and extruded at 100 nm. Plasma membrane vesicles used for comparison were produced as previously reported^31^. DNA functionalization of both viral particles and target liposomes was performed by adding DNA-lipids to either particles or liposomes at a concentration of 0.03 mol % lipid for liposomes and 0.2 μM for viral particles. The DNA sequences consist of a 24-mer for liposomes and a complementary 24-mer with a 24-mer poly-T spacer for viral particles; full sequences are given in the Supporting Information. These particular sequences have been extensively validated for tethering influenza viral particles in the past ^18^.

Biotinylated liposomes were bound to a PEGylated glass coverslip inside a microfluidic flow cell using PLL-PEG-biotin and neutravidin as previously described ^32^. DNA-functionalized viral particles were allowed to bind for 1-1.5 hours in the dark at room temperature, and then unbound virus was washed away, the flow cell chamber was brought to 37 °C, and the fusion reaction was initiated by addition of soluble protease at pH 7.4. Video micrographs were acquired via epifluorescence microscopy using a 100×, 1.49 NA oil immersion objective and an Andor Zyla 4.2 sCMOS camera. Micrographs were recorded at 1 s intervals using a 150-ms exposure time. Fluorescence time series data were analyzed using previously reported protocols ^18,31^ with Matlab code available from https://github.com/kassonlab/micrograph-spot-analysis. The number of fusion events compiled into each cumulative distribution function (CDF) is given in Table S1.

## RESULTS

We measured SARS-CoV-2 spike-mediated binding and fusion to both synthetic and cellular membranes, using synthetic liposomes to ensure the absence of endogenous ACE-2 receptor and employing Vero cell plasma membrane vesicles as a control. Experiments were performed using three SARS-CoV-2 spikes: Wuhan, D614G/N501Y and B.1.1.529 (Omicron) and using two viral particle types: pseudoviruses on an HIV core and virus-like particles created by co-expressing S, E, M, and N proteins. These were chosen to permit testing of SARS-CoV-2 entry requirements under BSL-2 conditions. In all cases, viral particles and target membranes were separately incubated with complementary DNA strands conjugated to DPPE lipids. Target membranes were then immobilized in a microfluidic flow cell, and viral particles were allowed to bind. After washing away unbound particles, soluble proteases were added to trigger fusion. In prior work, trypsin has been shown to activate SARS-CoV-2 S for fusion in the presence of ACE-2 receptor, with resulting fusion kinetics indistinguishable from S protein activated by either TMPRSS2 or cathepsin B or L ^31^. We thus use trypsin as the major protease in these DNA-tethering experiments, also presenting comparisons to TMPRSS2. Individual fusion events were monitored as a function of time since protease addition (Fig. 1).

**Figure 1.**
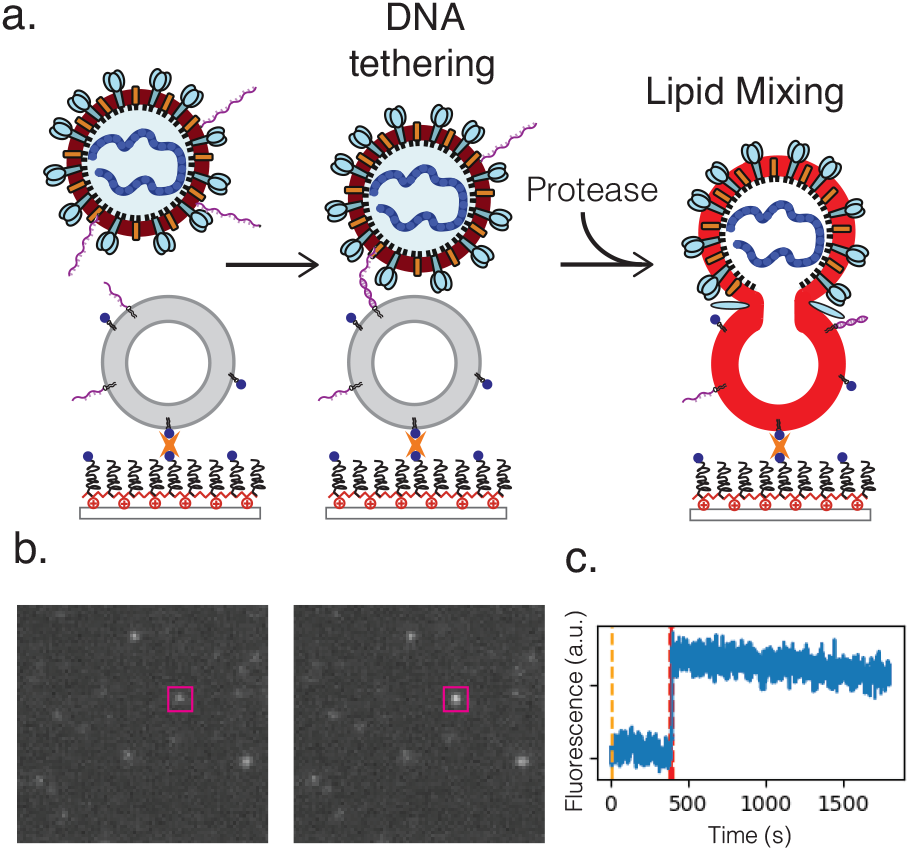
DNA-tethering and fusion of SARS-CoV-2 pseudoviruses and virus-like particles. The experiment design is schematized in panel (a), where DNA-functionalized viral particles are added to a microfluidic flow cell and allowed to bind to protein-free liposomes functionalized with complementary DNA. After unbound particles are washed away, fusion is initiated by addition of a soluble protease and monitored via lipid mixing, detected as fluorescence dequenching of Texas Red dye in the VLP or pseudoviral envelope. Representative images of a 10.1 × 9.6 μm sub-micrograph before and after lipid mixing are shown in panel (b) with a fusing particle outlined in magenta. The corresponding fluorescence intensity trace is plotted in panel (c).

Fusion events occurred only when viral particles were tethered to target membranes and only in the presence of protease. Fusion was primarily monitored via lipid mixing between the viral particle and the target membrane, detected via dequenching of Texas Red dye in the viral envelope and the resulting fluorescence enhancement. In prior work we have demonstrated that virus-like particles activated by exogenous protease can bind ACE-2 and productively enter cells in a manner that correlates with lipid mixing (even when endosomal acidification is inhibited)^31^. DNA-mediated viral binding was specific; when DNA was omitted, the number of adsorbed particles dropped >25-fold (Fig. S1). Similarly, fusion events were not observed in the absence of protease addition. Fig. 2 compares viral fusion kinetics for the three viral particle types used: Wuhan spike pseudotyped on an HIV core, Omicron (B.1.1.529) spike pseudotyped on an HIV core, and Wuhan D614G/N501Y virus-like particles. Both pseudoviruses tested yielded identical fusion rates, while the VLPs tested yielded rates that were slightly yet significantly faster (p ≤ 0.001, Kolmogorov-Smirnov test with Bonferroni correction for VLPs against all other samples shown in Figs. 2 and 3 except Omicron with 200 μg/mL trypsin; that sample yielded p = 0.04). This difference in fusion rate could result from differences in spike protein density on the virus-like particles versus the HIV core, but it could also be due to the presence of E and M in the virus-like particles, the percentage of active spike on the VLP surface, or the D614G/N501Y mutations. In particular, the D614G mutation has been previously characterized as favoring the “up” conformation of the spike receptor-binding domain^33–35^.

**Figure 2.**
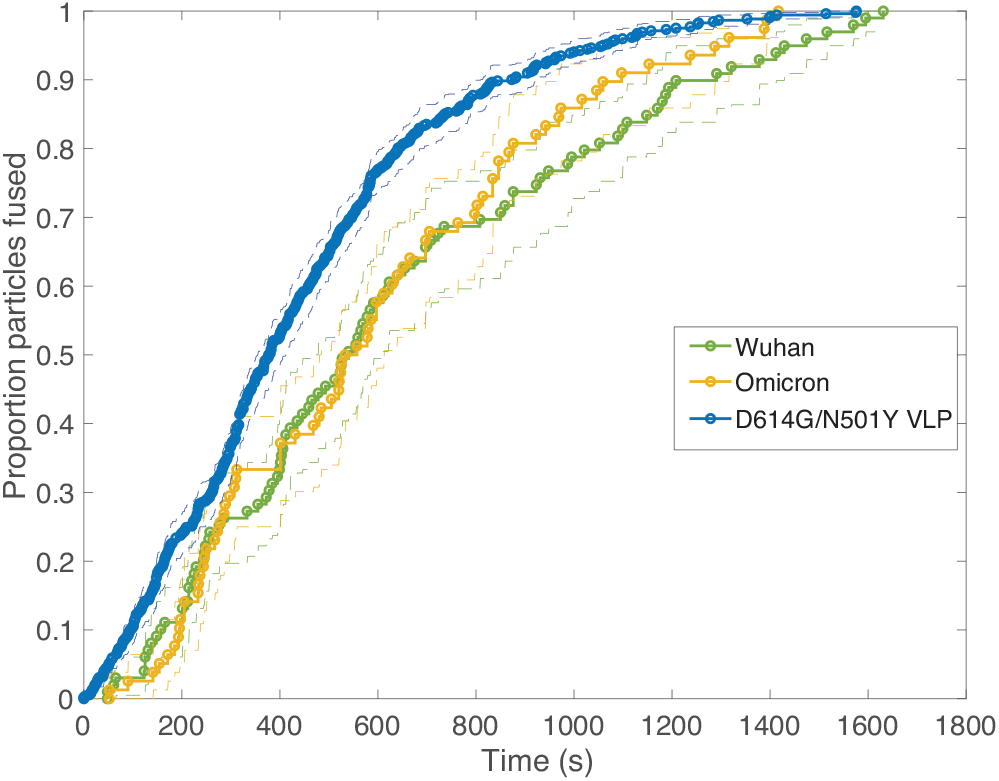
Cumulative distribution functions for fusion by different DNA-tethered SARS-CoV-2 spike particles. Lipid mixing is used as a surrogate for viral membrane fusion, and fusion kinetics are compared between HIV-based pseudoviruses displaying Wuhan or Omicron spike versus virus-like particles displaying D614G N501Y spike. Bootstrapped 90% confidence intervals are plotted in dashed lines. The D614G/N501Y virus-like particles fused significantly faster than any of the Wuhan or Omicron samples except Wuhan at 1000 μg/mL trypsin (p-values via 2-sample Kolmogorov Smirnov test of 0.001 for Omicron at 200 μg/mL, 0.04 for Wuhan at 200 μg/mL, 7×10-4 for Omicron at 500 μg/mL, 7×10-4 for Wuhan at 500 μg/mL, 0.002 for Omicron at 1000 μg/mL and 0.11 for Wuhan at 1000 μg/mL trypsin).

In order to probe the rate-limiting factors for SARS-CoV-2 fusion, we first varied the protease used for activation. In both the Wuhan and Omicron backgrounds, fusion kinetics were insensitive to trypsin concentration over a range from 200 to 1000 μg/mL (Fig. 3), and Wuhan was further tested and insensitive to trypsin concentration over a range from 10 to 1000 μg/mL (Fig. S2). TMPRSS2 did not efficiently activate Omicron spikes for fusion, consistent with prior reports^25,26^, but addition of 40 μg/mL soluble TMPRSS2 to D614G/N501Y virus-like particles yielded kinetics indistinguishable from 200 μg/mL trypsin (Fig. S3). These two results suggest that proteolytic cleavage is likely not the rate-limiting step for fusion of DNA-tethered viral particles, since changing protease concentration would be expected to alter enzyme-substrate complex formation and changing protease identity alters *k_cat_*.

**Figure 3.**
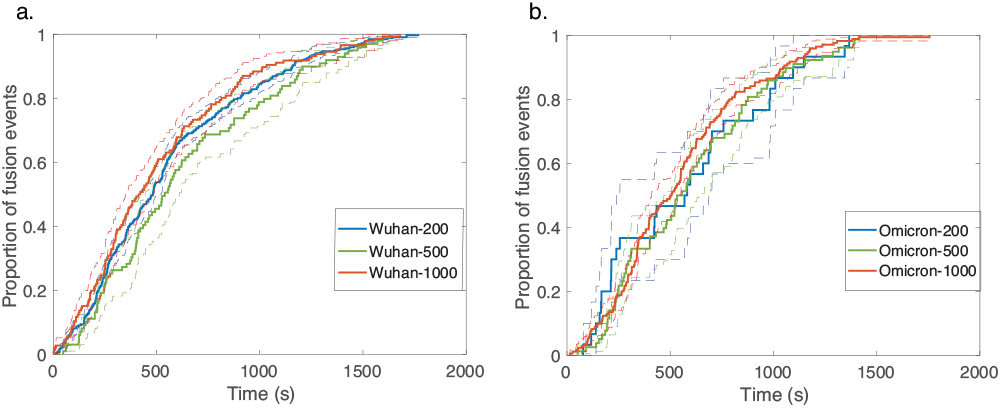
Cumulative distribution functions for fusion by DNA-tethered SARS-CoV-2 spike particles at different trypsin concentrations. Wu-han-pseudotyped particles are plotted in (a) at 200, 500, and 1000 μg/mL trypsin, and Omicron-pseudotyped particles are plotted in (b) over the same concentration range. Bootstrapped 90% confidence intervals are plotted in dashed lines. Wuhan and Omicron-pseudotyped particles fused at statistically indistinguishable rates at each trypsin concentration tested (p-values of 0.49 at 200 μg/mL, 0.84 at 500 μg/mL, and 0.09 at 1000 μg/mL, via 2-sample Kolmogorov Smirnov test) and similarly across trypsin concentrations for each of Omicron and Wuhan pseudovirus samples.

Since multiple spectroscopic and structural studies have suggested ACE-2 engagement may also alter SARS-CoV-2 S protein conformation^34,36–39^, we used soluble ACE-2 to test whether the protein can play an activating role separate from its attachment role. When soluble ACE-2 was added to the flow cell simultaneous to protease (after viral particle binding), we observed a significant speedup in lipid mixing kinetics with both the Wuhan and the Omicron spike proteins (Fig. 4). This suggests that the spike protein conformations required for fusion are accessible in the absence of ACE-2 yet are promoted by ACE-2 and indeed can be promoted by ACE-2 addition *in trans*. We also compared fusion driven by DNA tethering, trypsin activation, and soluble ACE-2 to “spontaneous” (i.e. ACE-2 and TMPRSS2) binding and fusion to the surface of Calu-3 plasma membrane vesicles. These results (Fig. 4c) show that the time between particle binding and fusion is faster than the time between protease introduction and fusion, suggesting that the 2D reaction kinetics and/or spatial organization of ACE-2 and TMPRSS2 on the plasma membrane surface may play a role. As a final comparison, we tested adding ACE-2 to the flow cell after DNA-binding but 15 minutes prior to protease addition. In this case, we observed 8-fold fewer fusion events (Fig. S4), suggesting that ACE-2 drives SARS-CoV-2 spike protein towards a conformational state that is fusion-enhanced but may also lead to inactivation if proteolytic cleavage does not occur. This is likely analogous to the well-known effect of pH on influenza hemagglutinin, where exposure to low pH activates hemagglutinin for fusion but also leads to inactivation over time if a target membrane is not present^40^.

**Figure 4.**
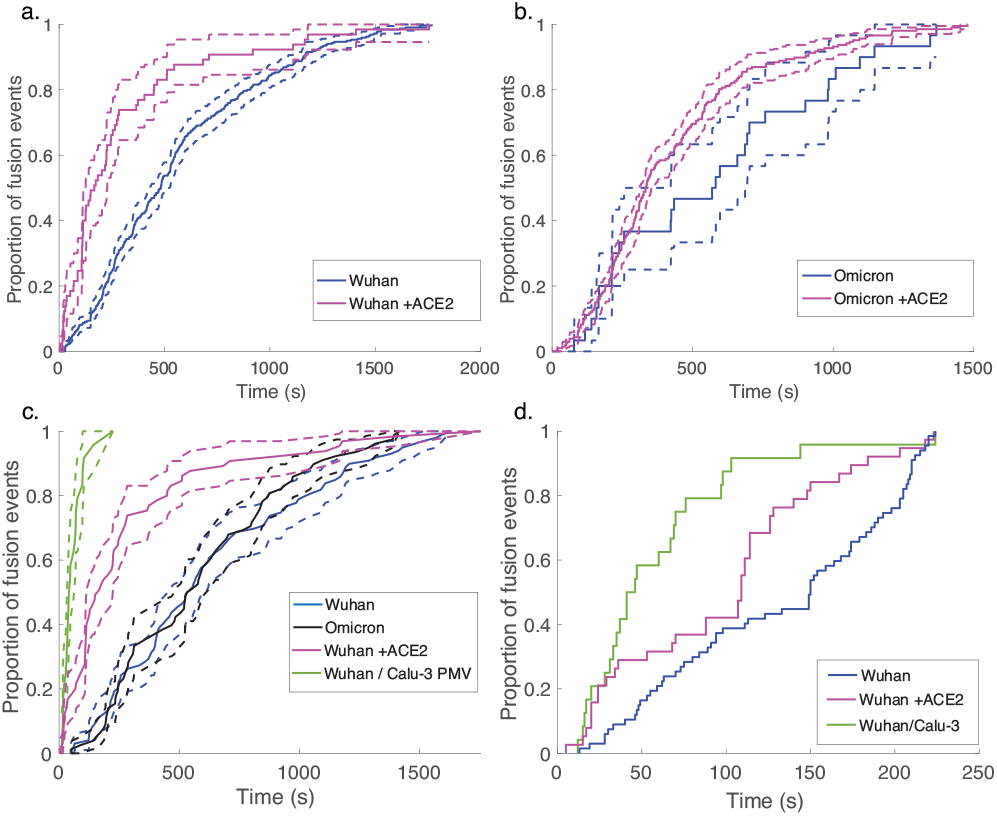
Addition of ACE-2 speeds fusion of DNA-tethered pseudovirions. Wuhan-pseudotyped particles are plotted in (a) with and without 40 μg/mL soluble human ACE-2, and Omicron-pseudotyped particles are plotted in (b). Rates of lipid mixing by DNA-tethered Wuhan pseudovirions with and without ACE-2 are compared to rates of lipid mixing by Wuhan pseudovirions to Calu-3 plasma membrane vesicles (PMV) containing both ACE-2 and TMPRSS2 in (c). Bootstrapped 90% confidence intervals are plotted in dashed lines. Because the longest waiting time measured for the plasma membrane vesicles was 224 s, cumulative distribution functions for the subset of DNA-tethered Wuhan pseudoviruses fusing in ≤ 224s were replotted in (d) to account for any potential sampling bias. When compared by either method, fusion to PMV was significantly faster (p < 0.001, 2-sample Kolmogorov Smirnov test).

Comparative analysis of the effect of ACE-2 on fusion mediated by the Wuhan versus the Omicron spike proteins shows a markedly greater enhancement for Wuhan versus Omicron. To further examine the source of this enhancement, gamma function fits were calculated for fusion by Wuhan and Omicron pseudoviruses in the absence and presence of ACE-2, fitting time-courses both independently and using a set of global parameters. The fraction of particles undergoing lipid mixing ***f*** is given by

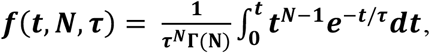

where Γ(N) is a gamma function. This functional form describes the idealized kinetics if fusion results from N events each with waiting time ***τ**^20,31^*, and the best global fits to the observed data were obtained for constant *τ* and variable N (Fig. S5). The addition of ACE-2 decreased N from 2.4 to 1.8 for Omicron and from 2.5 to 1.0 for Wuhan. Based on this, we conclude the most likely explanation for ACE-2’s action is that ACE-2 promotes a conformational change in the SARS-CoV-2 spike protein required for fusion. This change is one of ≥2 rate-limiting steps for fusion. The Wuhan spike is activated by soluble ACE-2 such that the ACE-2-related conformational change is no longer rate-limiting. For the Omicron spike, however, ACE-2 somewhat reduces the activation free energy, but this conformational change still contributes to the rate-limiting step. This model is consistent with other data suggesting that the Omicron spike is less conformationally labile than the Wuhan spike^41^. These results also suggests that the rate-limiting steps for SARS-CoV-2 fusion involve a stepwise process rather than a concerted mechanism.

## DISCUSSION

Using DNA-lipid tethers, we have shown that the ACE-2 receptor is not biochemically required for SARS-CoV-2 entry: if another factor can successfully bind the virus to a target membrane, protease activation can successfully trigger membrane fusion by the SARS-CoV-2 spike protein. However, when soluble ACE-2 is added *in trans*, it substantially speeds fusion. Together, these results yield a model where the protease-activatable conformations of SARS-CoV-2 are conformationally accessible in the absence of ACE-2 receptor but are substantially promoted by the presence of ACE-2. This is the functional equivalent of prior single-molecule FRET studies showing that RBD motions of the spike protein are accessible in the absence of ACE-2 but promoted by its presence^42^. Our data do not prove that the RBD motions correspond to the fusion-activatable conformations, but it is reasonable to hypothesize that they at least lead to such.

Kinetic analysis of both Wuhan and Omicron variant spike proteins in the absence and presence of ACE-2 suggest that ACE-2 activation drives one of the rate-limiting steps for fusion; in the presence of ACE-2, Wuhan displays only one kinetically evident rate-limiting step, while Omicron is less potently activated by ACE-2 and retains >1 rate-limiting step. This is consistent with the more “closed” conformational equilibria of Omicron and decreased ACE-2 responsiveness previously reported ^41^. It is satisfying that in analogy to many other viral spike proteins, ACE-2 activation in the absence of the factors required for full fusion can lead to spike inactivation. This is also consistent with structural data on S1 dissociation by liganded SARS-CoV-2 spike protein ^7^.

These results complement recent single-molecule FRET studies showing that the SARS-CoV-2 spike protein exists on the surface of lentiviral particles in a dynamic equilibrium between “closed” and “open” states, and that soluble ACE-2 promotes open conformations^42,43^. The precise relationship between the spike conformational equilibria and the pathway towards fusion remains incompletely determined, but the spectroscopic, structural, and now fusion kinetics data suggest that ACE-2 promotes conformational changes in the spike protein that are on-pathway for fusion. The single-molecule FRET data imply, and our data demonstrate, that the fusion-active conformations are energetically accessible in the absence of ACE-2, but that the addition of ACE-2 removes what is otherwise a rate-limiting free energy barrier on the fusion pathway.

The ability of SARS-CoV-2 spike to mediate VLP and HIV pseudovirus fusion to both simple synthetic liposomes and plasma membrane vesicles also provides an interesting counterpoint to observations of VSV pseudovirus trafficking in cells, where endocytosis appeared required for entry as well as mildly acidic pH^27^. It has long been observed that cell-cell fusion pathways mediated by SARS-CoV-2 spike as well as other viral fusion proteins can differ biochemically and mechanistically from virus-cell fusion^44,45^. Here, we show in a clean biochemical assay that relatively simple liposomes can support VLP fusion at neutral pH in the presence of an appropriate protease. The most likely entry pathway in cells may impose additional requirements, whether due to membrane residence time, increased efficiency of fusion in endocytic compartments, or other factors. This is why both observationally defining entry pathways in cells and biochemically defining requirements for fusion are critically important for understanding SARS-CoV-2 entry.

## CONCLUSIONS

The SARS-CoV-2 virus can utilize a variety of pathways for cell entry, depending both on cell type and viral variant. It is thus critical to determine the biochemical requirements for entry in order to ascertain both the mechanisms of viral fusion and the potential for future evolutionary plasticity. Here, we do so using DNA tethers as attachment factors in place of ACE-2. When so attached, both pseudoviruses and viruslike particles can fuse to synthetic liposomes, triggered by addition of a protease at neutral pH. ACE-2 is thus a high-affinity attachment factor but does not play a required role in conformational activation of the spike protein. By adding soluble ACE-2 *in trans*, however, we demonstrate that ACE-2 does promote fusion, speeding a required step in conformational activation so that it ceases to become rate-limiting in the Wuhan variant. Thus, a mechanistic picture of SARS-CoV-2 activation emerges where ACE-2 lies intermediate between influenza receptors which do not conformationally activate the viral glycoprotein and HIV receptors/co-receptors, which are typically required. This intermediate role permits SARS-CoV-2 to efficiently leverage ACE-2 but likely imparts a greater degree of evolutionary plasticity, since it has the potential to infect host cells using secondary attachment factors.

## Author Contributions

The manuscript was written through contributions of all authors. All authors have given approval to the final version of the manuscript.

## ACKNOWLEDGMENT

The authors thank Jesse Bloom, Jennifer Doudna, and Alejandro Balazs for the kind gift of plasmids. We also thank W. Mothes and R. Rawle for helpful discussions. This work was supported by grants from the Commonwealth Health Research Board (207-01-18), UVA Global Infectious Diseases Institute, and Knut and Alice Wallenberg Foundation (KAW2020.0209) to P.M.K.

## Notes

### Competing Interest Statement

The authors have declared no competing interest.

