## Supporting Information for "The ACE-2 receptor accelerates but is not biochemically required for SARS-CoV-2 membrane fusion"

Marcos Cervantes[1‡], Tobin Hess[1‡], Giorgio G. Morbioli[1‡†], Anjali Sengar[1], and Peter M. Kasson[1,2*]
1. Departments of Molecular Physiology and Biomedical Engineering, University of Virginia, Charlottesville VA 22908.
2. Science for Life Laboratory and Department of Molecular and Cellular Biology, Uppsala University, Uppsala SE 75123.

### Supporting Methods

#### *DNA-Lipids*

DNA-lipids were synthesized as previously described[1] using DPPE azide and DBCO-functionalized DNA. DPPE-TAG TAT TCA ACA TTT CCG TGT CGA was added to liposomes, and DPPE-TTT TTT TTT TTT TTT TTT TTT TTT TCG ACA CGG AAA TGT TGA ATA CTA was added to viral particles. The latter is complementary but includes a 24-mer poly-T spacer. These sequences correspond to sequences C and D previously reported for influenza tethering[1].

#### *Liposome Preparation*

Lipids (68.75 mol% POPC, 20% DOPE, 10 % chol, 1% biotin-DPPE, 0.25 % OG-DHPE) were mixed in chloroform and then dried under nitrogen to form a thin film. Films were stored under vacuum overnight, to ensure residual solvent removal, and then resuspended in 500 μL of reaction buffer (10 mM $NaH_2PO_4$, 90 mM sodium citrate, 150 mM NaCl, pH 7.4), and vortexed at low speeds for 5 min, until a milky suspension was formed. The suspension was subjected to 5 freeze-thaw cycles and then extruded 19 times through a 100 nm polycarbonate membrane, using a LiposoFast extruder (Avestin, Canada), to yield primarily large unilamellar vesicles. Lipid vesicles were stored in the dark, at 4 ºC, for no longer than 1 week. For DNA-tethering experiments, DNA-lipid conjugates were incubated with the liposome suspension overnight at 4 °C such that the final concentration of DNA-lipids was 0.03 mol % of lipids.

#### *Preparation of pseudoviruses and virus-like particles*

Pseudovirus production was performed in HEK 293T cells as previously described[2,3] using the following plasmids at ratios of 1: 0.22: 0.22: 0.22: 0.33: Luciferase-IRES-ZsGreen (BEI NR-52516) : HDM-Hgpm2 (BEI NR-52517) : pRC-CMV-Rev1b (BEI NR-52519) : HDM-tat1b (BEI NR-52518) : Spike- ALAYT (BEI NR-52515) following previously published protocols. All plasmids were gifts of Jesse Bloom. Pseudoviral supernatant was collected at 48 hours post transfection. Cellular debris was removed by centrifugation at 700 x g for 7 min at 4 ºC to remove cell debris followed by 0.45 μM filtration.

Virus-like particles were also prepared as previously described[4] by co-transfecting plasmids for N (Addgene 177937); M and E (Addgene 177938); and S (D614G N501Y; Addgene 177939) along with a luciferase gene with SARS-CoV-2 packaging sequence PS9 into 293T cells (Addgene 177942). Plasmids were gifts from Jennifer Doudna.

Supernatants collected above were pelleted through a 25% sucrose cushion in HEPES-MES buffer (20 mm HEPES, 20 mm MES, 130 mm NaCl (pH 7.4)) at 140,000 x g for 2 h, 4 °C. Pellets were resuspended in HEPES-MES (20 mM HEPES, 20 mM MES, 130 mM NaCl, pH 7.4) buffer without sucrose to obtain a 100x concentration of the initial volume. Viral aliquots were stored at -80 °C and were thawed no more than once. Pseudoviruses and VLPs were handled under BSL-2 conditions with institutionally approved safety protocols.

***Fluorescent labeling of pseudovirus particles and VLPs***

Pseudoviruses and VLPs were labeled with Texas Red-DHPE as follows. Briefly, 4 μL of TR-DHPE solution (0.75 mg/mL) were added to 240 μL of HEPES buffer. 200 μL of the staining solution were added to 50 μL of a 100X HIV pseudovirus or virus-like particle preparation (total viral protein concentration for HIV pseudovirus and virus-like particles were determined to be approximately 0.6 mg/mL and 2.9 mg/mL, respectively via a micro-BCA assay). After homogenization, the mixture was incubated in the dark on a rocker for 2 hours at room temperature. After the incubation, the solution was split into 2 centrifuge tubes, and 1.5 mL of HEPES buffer was added to each tube. Tubes were centrifuged for 1 hour at 4 °C, at 20000 rcf. The supernatant was carefully discarded, and the pellets were resuspended in 25 μL of HEPES buffer in each tube.

Viral particles were then functionalized with DNA lipids as follows. DNA lipids were added to Texas-Red-labeled viral particles at a final concentration of 0.4 μM and incubated overnight, in the dark, at 4 °C. Labeled pseudoviruses were stored in the dark at 4 °C for no longer than 1 week.

**Proteases and soluble ACE-2**

Trypsin and TMPRSS2 were each prepared as follows. TPCK-treated trypsin (Sigma-Aldrich) was dissolved in reaction buffer (10 mM $NaH_2PO_4$, 90 mM sodium citrate, 150 mM NaCl, pH 7.4), and TMPRSS2 (Creative Biomart) was dissolved in DI water. Both were kept as frozen aliquots at -20 °C at a concentration of 1,000 μg/mL, to be thawed, diluted, and warmed for immediate use for image acquisition.

ACE-2 (MP Biomedicals) was shipped in 8 M Urea, 20 mM Tris pH 8.0, 150 mM NaCl, 200 mM imidazole. Dialysis into 20 mM Tris pH 8.0, 150 mM NaCl, 200 mM imidazole was performed to remove urea using a dialysis slide with a 3.5 kDa cutoff (Thermo Scientific). The dialysis buffer was exchanged after 1.5 and 3 hours at room temperature; dialysis was continued overnight at 4 °C. The concentration of dialyzed ACE-2 was estimated at 812 μg/mL via absorbance at 280 nm. Aliquots were made and kept at -20 °C to be thawed and diluted for immediate use.

***Microfluidic Flow Cell Preparation***

Glass coverslips (24 x 40 mm, No 1.5, VWR International) were washed in a solution of 7x detergent (MP Biomedicals) and DI water at a ratio of 1:7 under continuous stirring and heating until the solution appeared clear, approximately 15 minutes. Detergent was removed by rinsing in excess volumes of DI water and coverslips were then annealed in a kiln for 4 hours at 400 °C. After cooling and several rinses with ultrapure water, coverslips were submerged in a bath sonicator for 5 minutes, rinsed with additional ultrapure water, sonicated again with absolute ethanol, and finally rinsed again with ultrapure water. Coverslips were then dried at 80 °C for several hours and stored in an airtight tube.

Microfluidic flow-cell molds were made using soft lithography as summarized below. Briefly, Kapton polyimide tape (Ted Pella) was laid on a glass surface where multiple flow channels (1 mm x 13 mm x 70μm) were cut using a Cameo 4 cutter-plotter (Silhouette) and excess tape was removed. Polydimethylsiloxane (PDMS; Sylgard 184) was poured into molds after mixing at a ratio of 10:1 elastomer : curing agent, degassed under vacuum and cured at 60 °C for 3 hours. After cooling, rectangular flow cells were sectioned out with influx/efflux holes made using a 2 mm biopsy punch.

Glass coverslips were plasma cleaned for 5 minutes (Harrick Plasma) and were then plasma bonded to PDMS flow cells after 1 minute plasma co-activation. Immediately afterwards, flow channels were flushed and coated with PLL-PEG : PLL-PEG-Biotin mixture (95% : 5%). After 30 minutes incubation at room temperature, channels were rinsed with 1 mL of ultrapure water and subsequently flushed with HEPES buffer (20 mM HEPES, 150 mM NaCl, pH 7.2). The channels were flushed with a 0.2 mg/mL solution of NeutrAvidin (Thermo Scientific), and the flow cell was incubated for 30 minutes at room temperature. After washing excess NeutrAvidin with 1 mL of HEPES buffer, channels were flushed with liposome reaction buffer (10 mM $NaH_2PO_4$, 90 mM sodium citrate, 150 mM NaCl, pH 7.4) and biotinylated liposomes functionalized with DNA-lipid conjugates were added, then left to incubate overnight at 4 °C. Channels were washed to remove excess liposomes with 1 mL of reaction buffer before addition of viral agents.

Texas-Red-labeled viral particles (either HIV-pseudovirus or virus-like particles) functionalized with DNA-lipid conjugates were diluted at an approximate ratio of 1:15 in 1.5% bovine serum albumin solution dissolved in liposome reaction buffer (10 mM $NaH_2PO_4$, 90 mM sodium citrate, 150 mM NaCl, pH 7.4) and immediately added to the flow cell channels. Flow cells were incubated at room temperature for 1 to 1.5 hours in darkness to allow for DNA-lipid conjugate binding and plastic reservoirs were affixed over the influx holes. Channels were rinsed with 1 mL of liposome reaction buffer to remove unbound viral particles, proteases and ACE-2 were diluted, added to reservoirs, and channels were imaged immediately at 37 °C.

**Fluorescence microscopy and image analysis**

Video micrographs were acquired using a Zeiss Axio Observer inverted microscope controlled using the MicroManager software[5]. The excitation light source was a Spectra X Light Engine (Lumencor), excitation/emission filters were 480/40 and 535/50 for Oregon Green and 560/40 and 630/75 for Texas Red. Images were acquired via a 100x oil immersion objective and a Zyla sCMOS 4.2 camera (Andor) at one-second intervals using a 150 ms exposure time. Micrographs were analyzed using previously reported single-virus detection and spot-tracking protocols,[6,7] with an additional manual review stage for fusion events. Matlab code is available from https://github.com/kassonlab/ micrograph-spot-analysis.

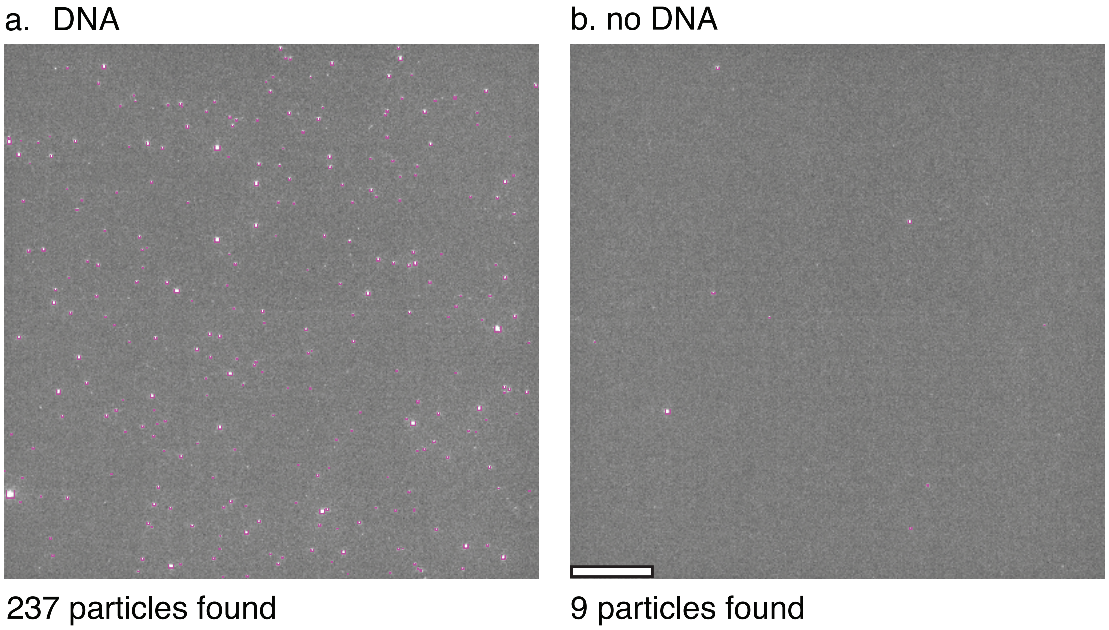


**Figure S1. Low binding of labeled viral particles in the absence of DNA tethers**. Micrographs show Texas-Red-labeled pseudovirus that was DNA-functionalized and allowed to bind to target liposomes either (a) containing complementary DNA-lipids or (b) without DNA-lipids. Unbound virus was washed away, and spots were counted, outlined in magenta. 237 particles bound when DNA-lipids were present, while only 9 particles bound without DNA-lipids. The latter likely represents nonspecific adsorption to the surface. Scale bar denotes 20 μm.

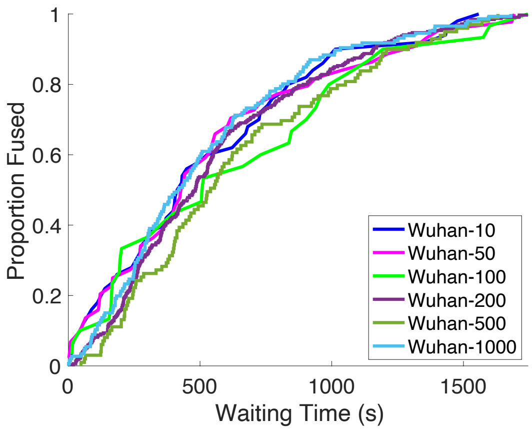


**Figure S2. Cumulative distribution functions for fusion by DNA-tethered Wuhan pseudoviruses at different trypsin concentrations.** Cumulative distribution functions are plotted for fusion experiments at ranges from 10 μg/mL to 1000 μg/mL trypsin. None of the distributions are significantly different via 2-sample Kolmogorov Smirnov tests (p-values range from 0.18 to 0.92, well above significance cutoffs when multiple hypothesis testing is accounted for), but all are significantly slower than when soluble ACE-2 is added to 200 μg/mL trypsin (all p-values < 0.0015 via 2-sample Kolmogorov-Smirnov tests).

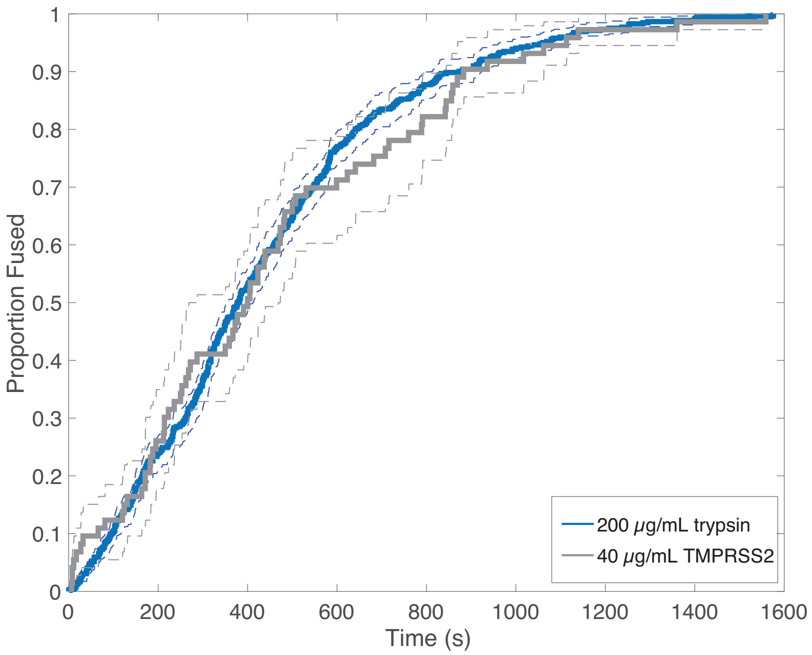


**Figure S3. Comparison of fusion kinetics for virus-like particles activated with trypsin and TMPRSS2.** The two distributions are statistically indistinguishable, with p > 0.73 via 2-sample Kolmogorov-Smirnov test.

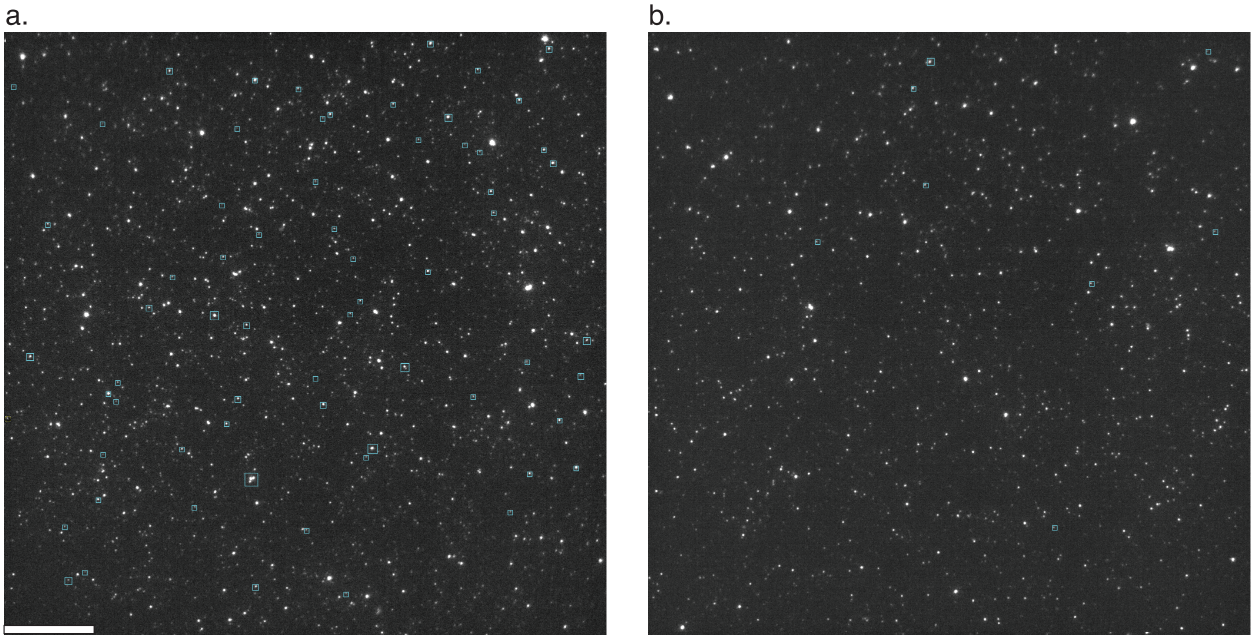


**Figure S4. Soluble ACE-2 enhances SARS-CoV-2 spike-mediated fusion when added simultaneous to protease but reduces when added prior**. Panels (a) and (b) show micrographs of Texas-Red-labeled pseudovirus tethered to liposomes after either addition of (a) 200 μg/mL trypsin simultaneous to soluble ACE-2 or (b) soluble ACE-2 added 15 minutes prior to trypsin. In panel (a), 65 dye dequenching events were observed, and the particles are outlined in cyan boxes. In panel (b), 8 dye dequenching events were observed. Scale bar denotes 20 μm.

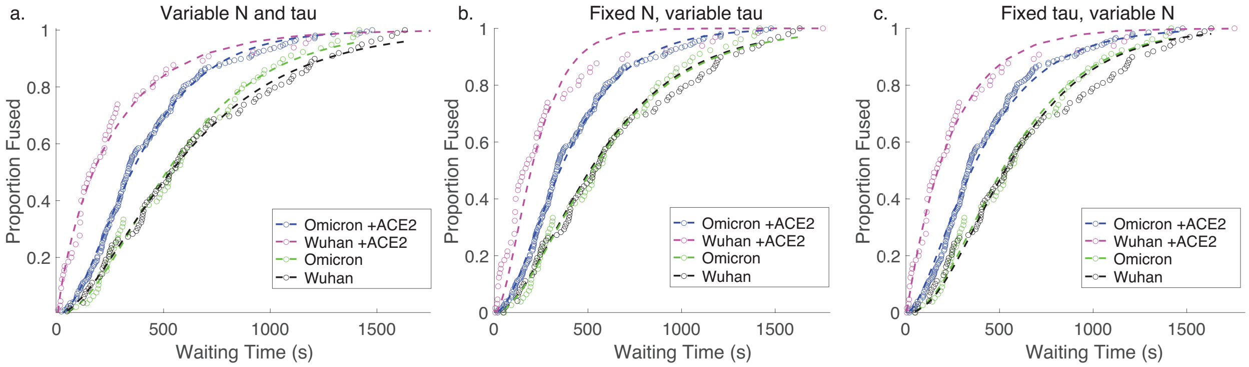


**Figure S5. Gamma-function fits to fusion kinetics.** Gamma-function fits were calculated for lipid-mixing CDFS of Omicron and Wuhan pseudoviruses with and without ACE-2. These fits were calculated either (a) varying N and tau independently, (b) fixing N globally across the data sets and varying tau, or (c) fixing tau and varying N.

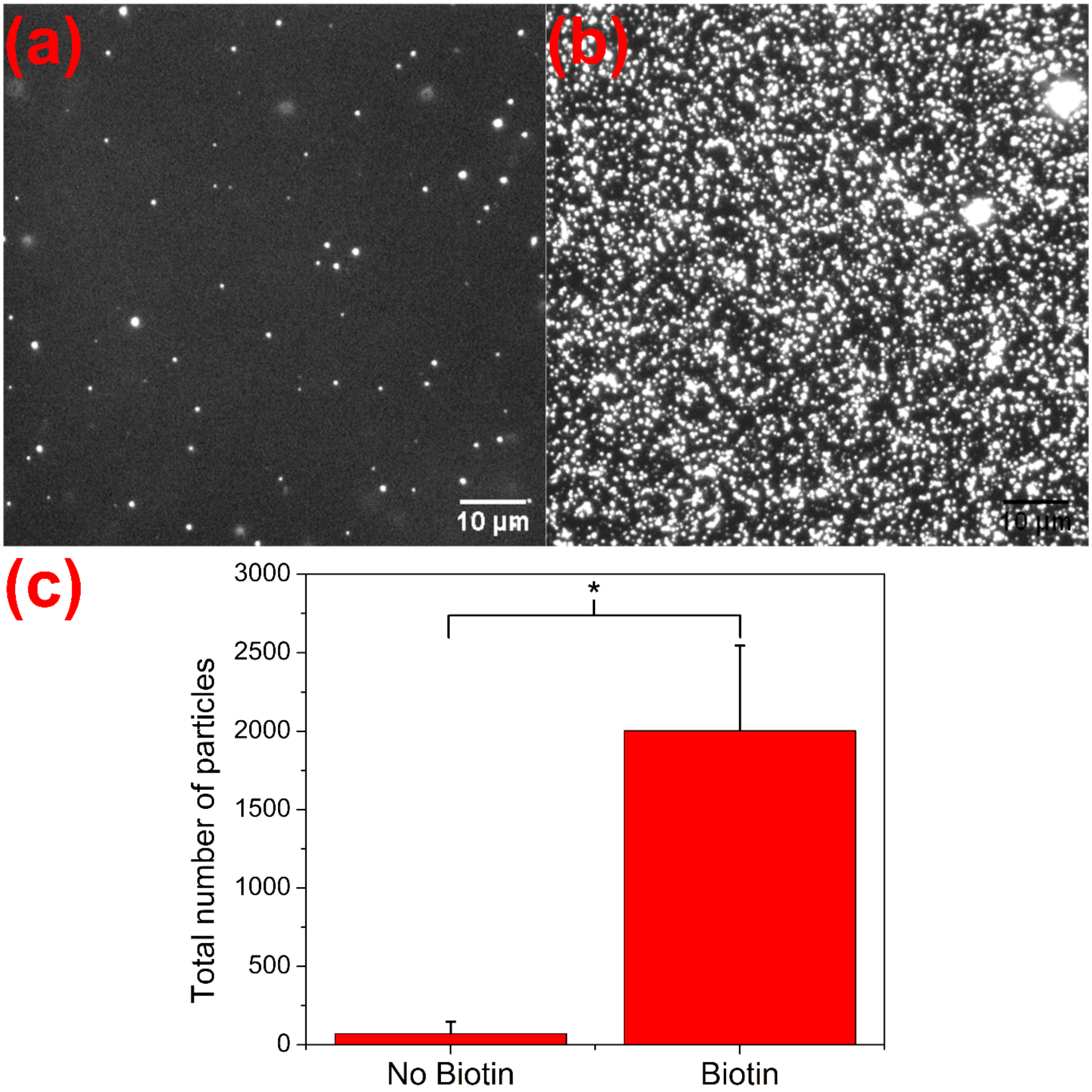


**Figure S5. Low nonspecific binding in the absence of biotinylation.** Micrographs in panels (a) and (b) show Oregon Green-labeled liposomes bound to the flow cell surface in (a) the absence of PLL-PEG-Biotin and (b) the presence of 2.5 % PLL-PEG-Biotin and neutravidin. Liposomes bonded to the flow cell channel surface via biotin-neutravidin-biotin binding. The number of bound liposomes is plotted in panel (c). In the presence of biotin, >20x more liposomes bound, $p < 0.05$ via t-test. Scale bars in the micrographs denote 10 μm. Micrographs were recorded using an Andor iXon Ultra 897 EMCCD camera.

| **Pseudovirus:** | **Protease:** | **Co-factor:** | **# of lipid mixing events:** |
|---|---|---|---|
| Wuhan / HIV pseudovirus | 10 μg/mL trypsin | | 52 |
| Wuhan / HIV pseudovirus | 50 μg/mL trypsin | | 44 |

| | | | |
|---|---|---|---|
| Wuhan / HIV pseudovirus | 100 µg/mL trypsin | | 30 |
| Wuhan / HIV pseudovirus | 200 µg/mL trypsin | | 320 |
| Wuhan / HIV pseudovirus | 500 µg/mL trypsin | | 99 |
| Wuhan / HIV pseudovirus | 1000 µg/mL trypsin | | 146 |
| Wuhan / HIV pseudovirus | 200 µg/mL trypsin | ACE-2 | 65 |
| Omicron / HIV pseudovirus | 200 µg/mL trypsin | | 30 |
| Omicron / HIV pseudovirus | 500 µg/mL trypsin | | 78 |
| Omicron / HIV pseudovirus | 1000 µg/mL trypsin | | 170 |
| Omicron / HIV pseudovirus | 200 µg/mL trypsin | ACE-2 | 207 |
| D614G, N501Y / VLP | 200 µg/mL trypsin | | 522 |
| D614G, N501Y / VLP | 40 µg/mL TMPRSS2 | | 73 |
| Wuhan / HIV pseudovirus | | PMV containing ACE-2 and TMPRSS2 | 24 |

**Table S1. Numbers of fusion events observed for each condition reported.**
